# Visualizing Wnt secretion from Endoplasmic Reticulum to Filopodia

**DOI:** 10.1101/271684

**Authors:** Naushad Moti, Jia Yu, Gaelle Boncompain, Franck Perez, David M Virshup

## Abstract

Wnts are a family of secreted palmitoleated glycoproteins that play a key role in cell to cell communications during development and regulate stem cell compartments in adults. Wnt receptors, downstream signaling cascades and target pathways have been extensively studied while less is known about how Wnts are secreted and move from producing cells to receiving cells. We used the synchronization system called Retention Using Selective Hook (RUSH) to study Wnt trafficking from endoplasmic reticulum to Golgi and then to plasma membrane and filopodia in real time. Consistent with prior studies, inhibition of porcupine (PORCN) or knockout of Wntless (WLS) blocked Wnt exit from the ER. Indeed, WLS was rate-limiting for Wnt ER exit. Wnt-containing vesicles paused at sub-cortical regions of the plasma membrane before exiting the cell. Wnt-containing vesicles were transported to adjacent cells associated with filopodia. Increasing the number of filopodia by expression of LGR5 in the producing cell increased the ability of a cell to send a Wnt signal. The RUSH system is a powerful tool to provide new insights into the Wnt secretory pathway.

## Introduction

Wnt proteins are secreted morphogens that play an important role in a variety of biological processes ranging from embryonic development, proliferation, differentiation, adult tissue homeostasis and cancer (Anastas & Moon 2013; Clevers & Nusse 2012; Yu et al. 2014). Wnts bind to cell surface receptors to activate diverse signaling pathways, the best-studied of which leads to the stabilization of β-catenin and the activation of target gene expression. Less is known about how Wnts travel from one cell to engage receptors on neighboring cells (Yamamoto et al. 2013; Yu & Virshup 2014; Stanganello & Scholpp 2016).

Newly synthesized Wnts are targeted to the lumen of the endoplasmic reticulum (ER) by signal peptides. There, they are modified by addition of a single mono-unsaturated palmitoleate by the acyl transferase PORCN (porcupine) (Krausova et al. 2011; Takada et al. 2006; Yu et al. 2014; Yu & Virshup 2014). Palmitoleation is required for the next step, binding to the integral membrane carrier protein Wntless (WLS) in the ER (Coombs et al. 2010; Yu et al. 2014; Herr & Basler 2012). The Wnt-WLS complex then shuttles to the Golgi where additional posttranslational modifications of Wnt can occur (Komekado et al. 2007). From the Golgi, Wnts bound to WLS move to the plasma membrane and then on to a Wnt receiving cell (Bänziger et al. 2006; Bartscherer et al. 2006; Herr & Basler 2012).

Several contrasting models have been proposed to explain how Wnts travel from the producing cell to the receiving cell (reviewed in Stanganello & Scholpp 2016). All models must explain how the hydrophobic palmitoleate moiety on Wnt is protected from an aqueous environment. Similarly, they must explain how Wnts can be transferred across basement membranes, e.g. in the intestine where Wnts produced in stromal cells regulate crypt base columnar cells (Kabiri et al. 2014; Krausova et al. 2011; Greicius et al. 2018). The simplest model is diffusion, where Wnts transfer in a gradient dependent manner, perhaps interacting with soluble Wnt binding proteins or with glycoproteins on the cell surface (Franch-Marro et al. 2005). Wnts have also been identified on exosomal vesicles in both Drosophila and mammalian systems (Gross et al. 2012; Stanganello et al. 2015; Saha et al. 2016). Another means by which Wnt transportation occurs is through WLS-containing exosomes across synapses, demonstrated in *Drosophila* (Korkut et al. 2009; Koles et al. 2012; Gross et al. 2012). Finally, Wnts may travel on signaling filopodia, also called cytonemes (Sagar et al. 2015; Korkut et al. 2009; Huang & Kornberg 2015). Indeed, in zebrafish models, filopodia have been visualized transporting Wnt8A from one cell to another (Stanganello et al. 2015).

The variety of potential mechanisms suggests there is a need to develop new experimental approaches to visualize Wnts during secretion. Previous studies have shown that there are several routes for protein secretion and transport in the cells. In order to dissect the route that a particular cargo protein uses, specialized assays are needed that can be adaptable to a variety of cargo proteins present in a cell (Boncompain et al. 2012). The temperature block assay that retains protein in the ER at 15 °C and in the Golgi at 20 °C (Matlin & Simons 1983; Saraste & Kuismanen 1984), has been a powerful tool to study intracellular trafficking of various proteins in living cells but these assays are limited and do allow to study cells in physiological conditions. The reasons for such blocks are still elusive. A method that relies on retention-release mechanism of cargo molecules was developed a few years ago (Boncompain et al. 2012). The system, called Retention Using Selective Hook (RUSH), contains two components. The protein of interest is fused to the streptavidin-binding peptide (SBP) and a genetically encoded fluorescent reporter (Boncompain et al. 2012). Second, streptavidin is fused to a protein stably located in the ER - this can be as simple as a KDEL sequence - and thus serves as the ER hook. At steady state, streptavidin-KDEL will be bound to SBP and the cargo will be retained in the ER. Upon biotin addition, cargo proteins with their fluorescent tags will be released from the streptavidin-KDEL. Their subsequent trafficking can be visualized using real-time live cell imaging in living cells (Boncompain et al. 2012) (Figure 1a).

**Figure 1:**
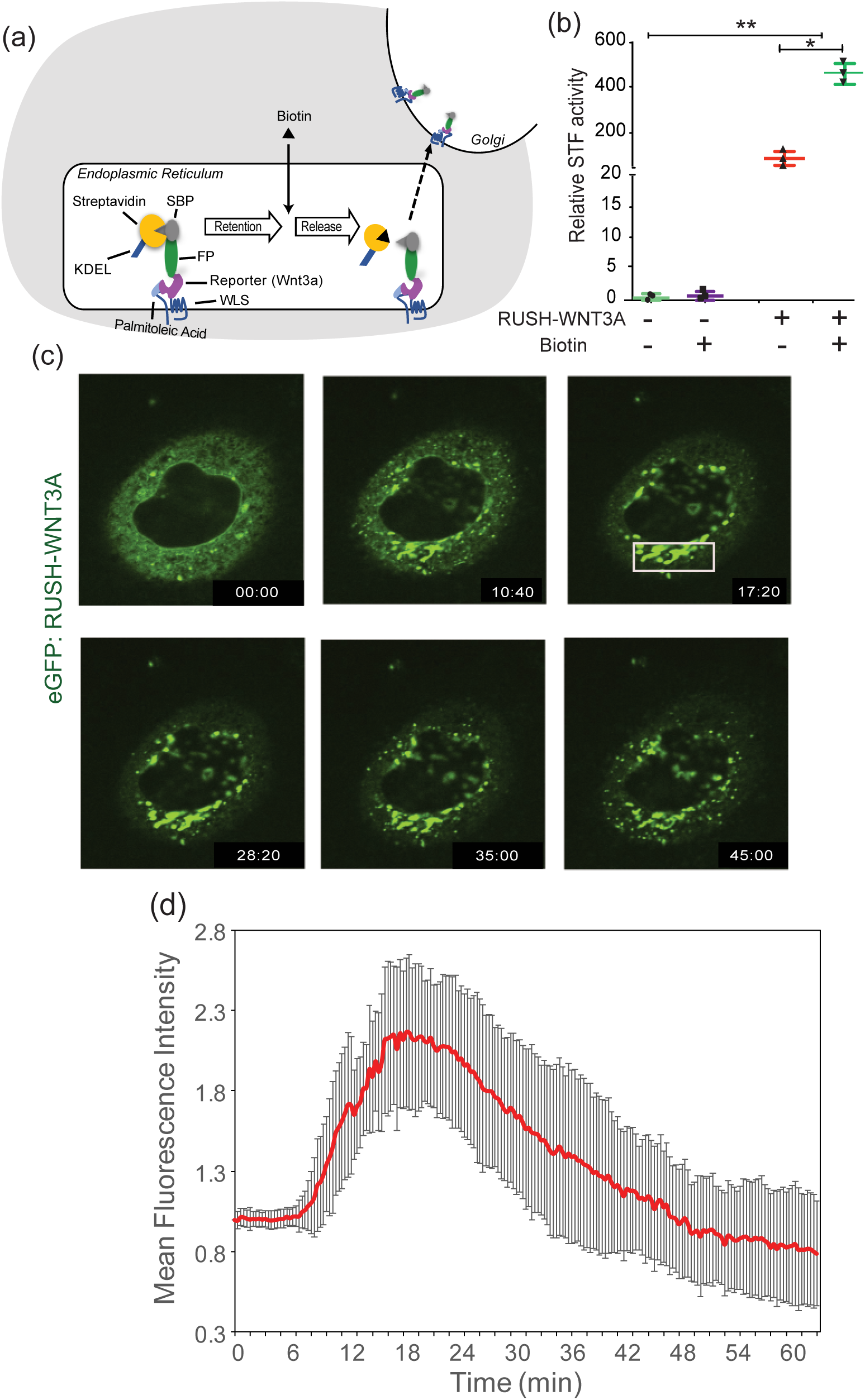
Synchronization of WNT3A protein using RUSH system. (a) Schematic of the RUSH-Wnt system. Streptavidin-KDEL interacts with SBP-GFPWnt to retain the protein in the ER. Wnt release is induced by the addition of biotin. (b) RUSH-WNT3A is signaling-competent. β-catenin-dependent luciferase expression was stimulated by transient expression of the indicated proteins in SuperTopFlash cells. 24 hours after transfection, luciferase activity was measured with and without biotin treatment. (\*\**p*<0.01; \**p*<0.05) (c) Indirect immunofluorescence photomicrographs of HeLa cells expressing RUSH-WNT3A at various time points after biotin addition, from Supplemental Video 1A. At time 00:00 (minutes:seconds) of biotin addition, WNT3A remained in the ER. At time 10:40, WNT3A can be seen in the Golgi. At time 17:20, WNT3A is seen entirely in Golgi. Starting around 28:20 and continuing at 35:00 WNT3A can be seen in vesicle exiting the Golgi and moving towards PM. By time 45:00, the Golgi begins to be depleted of WNT3A. (d) Time-dependent analysis of Golgi localization of RUSH-WNT3A. The plot shows fluorescence intensity in the Golgi region (white box in Figure (c)) at different time point after biotin addition. Intensities were normalized to the initial Golgi intensity with arbitrary units. (n=7 independent cells ± SD)

Here, using RUSH, we followed the path of Wnt proteins from the ER to the membrane. The studies confirm core insights regarding Wnt secretion and provides new evidence for the involvement of filopodia in transmitting Wnt signals. The RUSH-Wnt model will be useful system to study post-ER regulators of Wnt transport machinery and may be amenable to high-content screening for novel Wnt therapeutic targets.

## Material and Methods

### Cells and Reagents

HeLa, HEK293T and RKO cells were cultured in Dulbecco's modified Eagle medium (DMEM) (Invitrogen) supplemented with 10% of Fetal Bovine Serum (FBS) (Hyclone), 1% sodium pyruvate (Lonza) and 1% penicillin and streptomycin (Invitrogen) (complete medium) at 37 °C and 5% of CO_2_. HeLa and RKO cells were transfected using Lipofectamine 2000 Reagent (Invitrogen) using manufacturer’s protocol. Biotin was purchased from Sigma and used at a concentration of 100 μM. CellMask™ Deep Blue and Deep Green plasma membrane dyes were purchased from Invitrogen. The PORCN inhibitor ETC-159 was formulated in DMSO and used at a final concentration of 100 nM as previously described (Madan et al.).

### Plasmids

The RUSH plasmid, streptavidin-KDEL as a hook and SBP-eGFP/mCherry-E-cadherin as a reporter has been described previously (Boncompain et al. 2012). Briefly, E-cadherin was removed by restriction digestion using *FseI* and *SfiI* enzymes. *WNT3A* and *WNT8A* (Najdi et al.) were cloned by PCR. Plasmids Str-KDEL_SBP-mCherry-WNT3A and Str-KDEL_SBP-EGFP-WNT3A will be deposited at Addgene.

The STF reporter cell line (HEK293T cells with the firefly luciferase gene under the control of eight tandem repeats of TOPFlash TCF/LEF responsive promoter element) was a gift from Kang Zhang (University of California San Diego, La Jolla, CA) (Xu et al. 2004).

### CRISPR-Cas9

The WLS CRISPR knockout used pLentiCRISPRv2-Cas9 from Zhang’s lab (Addgene plasmid #52961) (Sanjana et al. 2014) with guide RNAs (gRNAs). Non-target control gRNA targeting sequences was GTTCGACTCGCGTGACCGTA, and WLS gRNA targeting sequences was AGGGGGGGCGCAAAAATGGC. For the knockout, RKO cells were transfected with 2 μg of either non-target control gRNA or WLS gRNA in a 60-mm dish and selected using puromycin. Minimal cell dilution was used to isolate single clones. The knockout clones were verified by both qPCR and western blotting.

### Live cell imaging and Image quantification

HeLa cells were seeded onto Nunc^TM^ Lab-Tek^TM^ 4-well chambered cover glass with non-removable wells (ThermoFisher Scientific) one day prior to transfection. 24 hours after transfection with RUSH-WNT3A expression plasmids, the medium was removed and D-biotin (Sigma-Aldrich) at 100 μM final concentration was introduced in the chamber. Time-lapse acquisition and z-stack imaging was done at 37 °C in a thermostat-controlled stage-top incubator using N-STORM with Andor CSU-W1 spinning disk confocal microscope (Nikon). Images were acquired using a 60x objective and NIS-Element Advanced Research software (Nikon). Raw images were processed using Imaris 9.0 (Bitplane, Oxford Instruments) and analysed using Fiji 2.0, an image processing package of ImageJ (Schindelin et al. 2012). FiloQuant, a user friendly ImageJ plugin, was used to detect and quantify filopodia (Jacquemet et al. 2017).

### STF Wnt signaling assay

HEK293T STF cells were seeded on 24-well plate and transfected with RUSH-WNT3A expression plasmids. 6-8 hours later, 100 μM biotin was added. After an additional 18 hours, transfected cells were washed with PBS and then lysed with 100 μl per well of 1x Reporter Lysis Buffer (Promega) supplemented with protease inhibitor cocktail (Roche). 20 μl of cell lysate was mixed with 50 μl of luciferase assay reagent (Promega), and a 1 sec integration time was used for measurement. Luminescence data were obtained using a Tecan Infinite M211 microplate reader and normalized to lactate dehydrogenase (LDH) as described previously (Coombs et al. 2010). GraphPad Prism 5.0 was used to quantify the data.

### Flow Cytometry

HeLa cells were seeded on 24-well plates and transfected with RUSH-*WNT3A* plasmid. After 24 hours, 100 μM biotin was added with or without 100 nM ETC-159. The cells were collected at different time points by trypsinizing with 1x PBS with 0.5 mM EDTA. The detached cells were washed twice with ice-cold 1x PBS + 1% FBS and incubated with anti-mCherry AlexaFluor 488 (ThermoFisher Scientific) conjugated antibody for 45 min. The cells were washed twice with ice-cold 1x PBS + 1% FBS and events were captured with MACSQuant^®^ VYB (Miltenyi Biotec). A total of 25,000 events were acquired and the data analyzed using FlowJo™ 10.

## Results

### Synchronization of WNT3A protein secretion using the RUSH system

To study the Wnt secretory and transport mechanism, we made use of the robust RUSH (Retention Using Selective Hook) system (Boncompain et al. 2012), which allows to analyze protein both qualitatively and quantitatively in real-time, and adapted WNT3A to the RUSH (Figure 1a). We first tested if the RUSH-Wnt fusion constructs retained biological activity. A SBP-GFP-WNT3A construct expressed in HEK293 cells with an integrated SuperTopFlash reporter (Xu et al. 2004) produced low levels of Wnt/β-catenin signaling, while the addition of biotin to the culture medium increased signaling 20-fold (Figure 1b), demonstrating that the fusion protein was competent to signal. We next followed the secretion of Wnt protein. We used HeLa cells because their morphology in culture made it easier to visualize the various cellular compartments. After expression in HeLa cells, SBP-GFP-WNT3A accumulate in the ER (Figure 1c, Supplementary video S1A). Upon biotin addition at time=00:00 (minutes:seconds), WNT3A was transported from ER to Golgi in a time dependent manner (Supplementary Video S1A; Figure 1c). At t=17:20, most of the WNT3A was transported to Golgi (Figure 1c). Wnt protein resided in the Golgi for approximately 20 minutes, presumably transiting from cis to trans-Golgi compartments and undergoing glycosylation. Subsequently, WNT3A was transported to the PM in small vesicles (Figure 1c). This pathway was not specific to WNT3A, as a RUSH-WNT8A construct showed similar results (Supplementary Figure S1; Supplementary Video S1B).

### Wnt exit from ER requires palmitoleation and WLS

In order to further assess if the RUSH-WNT3A constructs followed the known Wnt secretion pathway, we examined the effect of inhibited PORCN and WLS activity on eGFP-Wnt secretion. PORCN palmitoleates Wnt proteins in the ER, and loss of PORCN activity blocks Wnt secretion and activity (Madan et al. 2016; Tanaka et al. 2000; Zhai et al. 2004). Inhibition of PORCN activity with the PORCN inhibitor ETC-159 abrogated signaling from the eGFP-RUSH-WNT3A fusion protein, confirming the fusion protein behaves like native WNT proteins (Figure 2a). Live cell imaging shows that the same concentration (100 nM) of ETC-159 abolished WNT3A exit from the ER (Figure 2b, 2c; Supplementary Video S2). Figure 2d shows quantification normalized to Golgi intensity in Figure 2b (white box).

**Figure 2:**
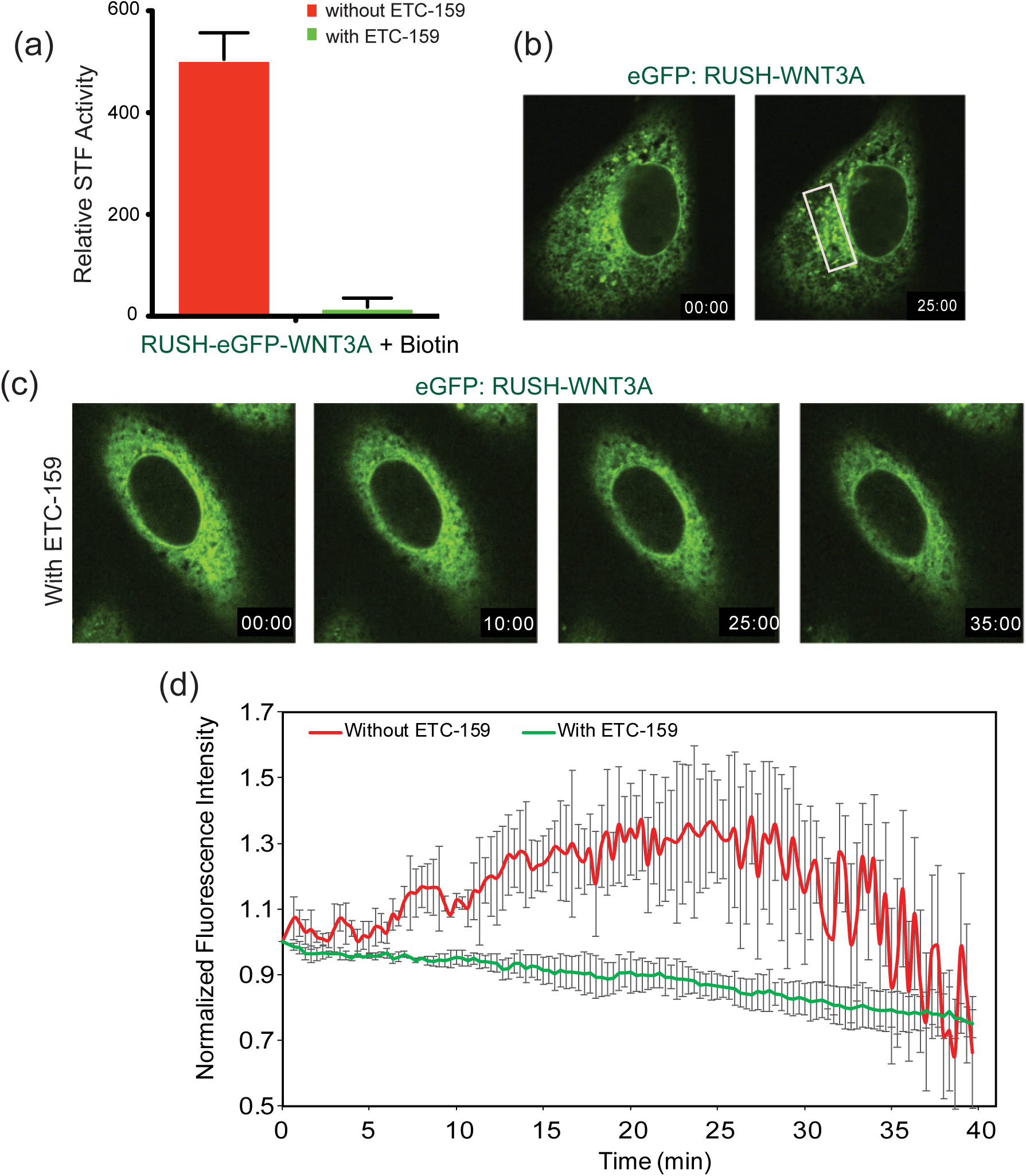
PORCN inhibition traps WNT3A in the ER. (a) SuperTopFlash assay on HEK293 cells with and without ETC-159. Luciferase readings taken 24 hours post-transfection with and without ETC-159 shows attenuated signaling upon PORCN inhibition. (b-c) Time course of WNT3A trafficking as in Fig 1, except that 100 nM ETC-159 was added to the cells 18 hours prior to the addition of biotin where indicated. (d) Time-dependent analysis of Golgi localization of RUSH-WNT3A in HeLa cells treated with and without ETC-159, as in Fig. 1. (n=3 cells per condition ± SD)

Following palmitoleation, Wnts bind to WLS in the ER and are transported to the Golgi. To further understand the role of WLS in Wnt trafficking and signaling, we employed the CRISPR-Cas9 system to knockout *WLS* in RKO cell lines that are normally WLS-high. Figure 3a shows that we achieved knockout of WLS at the protein level in two independent cell lines. Confirming the requirement for WLS, both knockout lines lost Wnt/β-catenin signaling and could be rescued by re-expression of wildtype WLS (Figure 3b). Live cell imaging of cells transfected with RUSH-WNT3A showed Wnt exit from the ER was blocked in RKO WLS KO cells (Figure 3c, 3d; Supplementary Video S3A). Re-expression of WLS in the RKO WLS KO cells rescued the trafficking, allowing WNT3A to move out of the ER (Supplementary Video S3B). Figure 3e quantitates the time course of Wnt ER to Golgi trafficking. These data illustrate the important roles of PORCN and WLS in WNT3A trafficking and further confirm the physiological nature of the RUSH-WNT3A construct.

**Figure 3:**
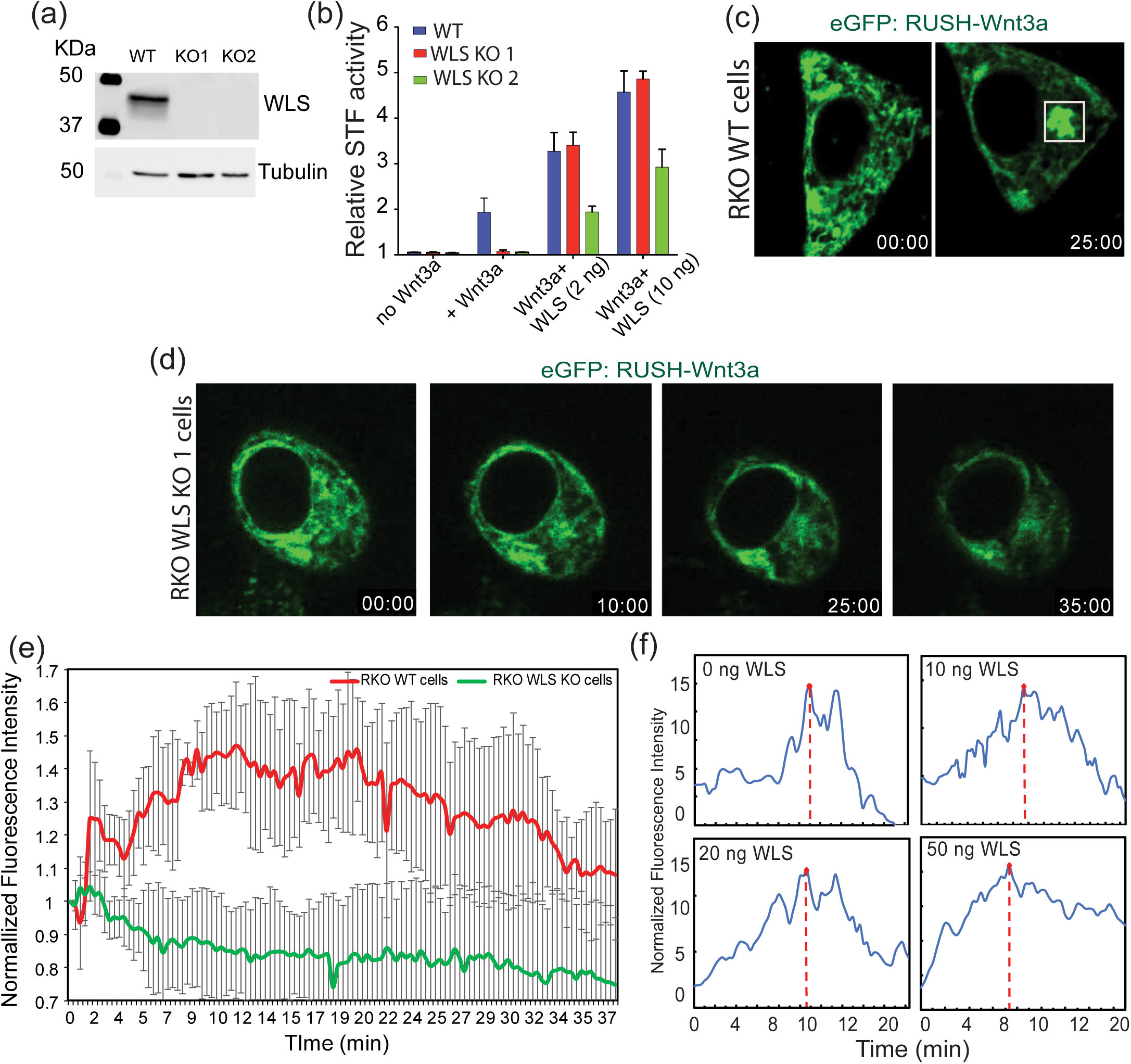
WLS knockdown abolishes WNT3A trafficking. (a) Western blotting shows complete loss of WLS protein in CRISPR-Cas9 *WLS* KO RKO cells. (b) SuperTopFlash assay shows complete loss of Wnt signaling in *WLS* KO cells and rescue by wildtype WLS re-expression. (c) Micrographs of wildtype RKO cells transfected with RUSH-WNT3A showing WNT3A moves from ER (time 00:00) (minutes:seconds) to Golgi (time 25:00) after biotin addition. From Supplemental Video 3A (d) WNT3A does not exit the ER in *WLS* KO cells. (e) Time-dependent analysis of ER-Golgi localization of RUSH-WNT3A in RKO wildtype and WLS KO RKO cells, as above. (n=4 cells per condition ± SD) (f) Increasing WLS abundance increases rate of WNT3A exit from the ER. Time-dependent analysis of Golgi localization of RUSH-WNT3A in HeLa cells (n=5 cells per condition)

We quantified the accumulation of WNT3A in the Golgi over time after biotin addition as in Figure 1d in HeLa cells. The transit time from ER to Golgi was longer than seen with other cargos such as RUSH-E-cadherin and RUSH-MannII (Boncompain et al. 2012). Wnt secretion is unique in that it requires both palmitoleation and subsequent binding to the carrier protein WLS to exit the ER (Yu et al. 2014). Hence, we tested if WLS was rate-limiting for WNT-ER exit. Indeed, co-expression of increasing amounts of WLS shortened the time of arrival of WNT3A at the Golgi (Figure 3f). Hence, WLS abundance is rate-limiting for Wnt ER exit.

### Wnt vesicles pause in sub-membranous region

We next followed the path of Wnt from Golgi to plasma membrane. Live cell z-stack imaging of HeLa cells expressing RUSH-WNT3A revealed WNT3A vesicles pausing immediately below the plasma membrane for up to several minutes (PM) before disappearing at the PM (Figure 4a-f; Supplementary Video S4). We did not observe re-uptake of RUSH-WNT by endocytosis.

**Figure 4:**
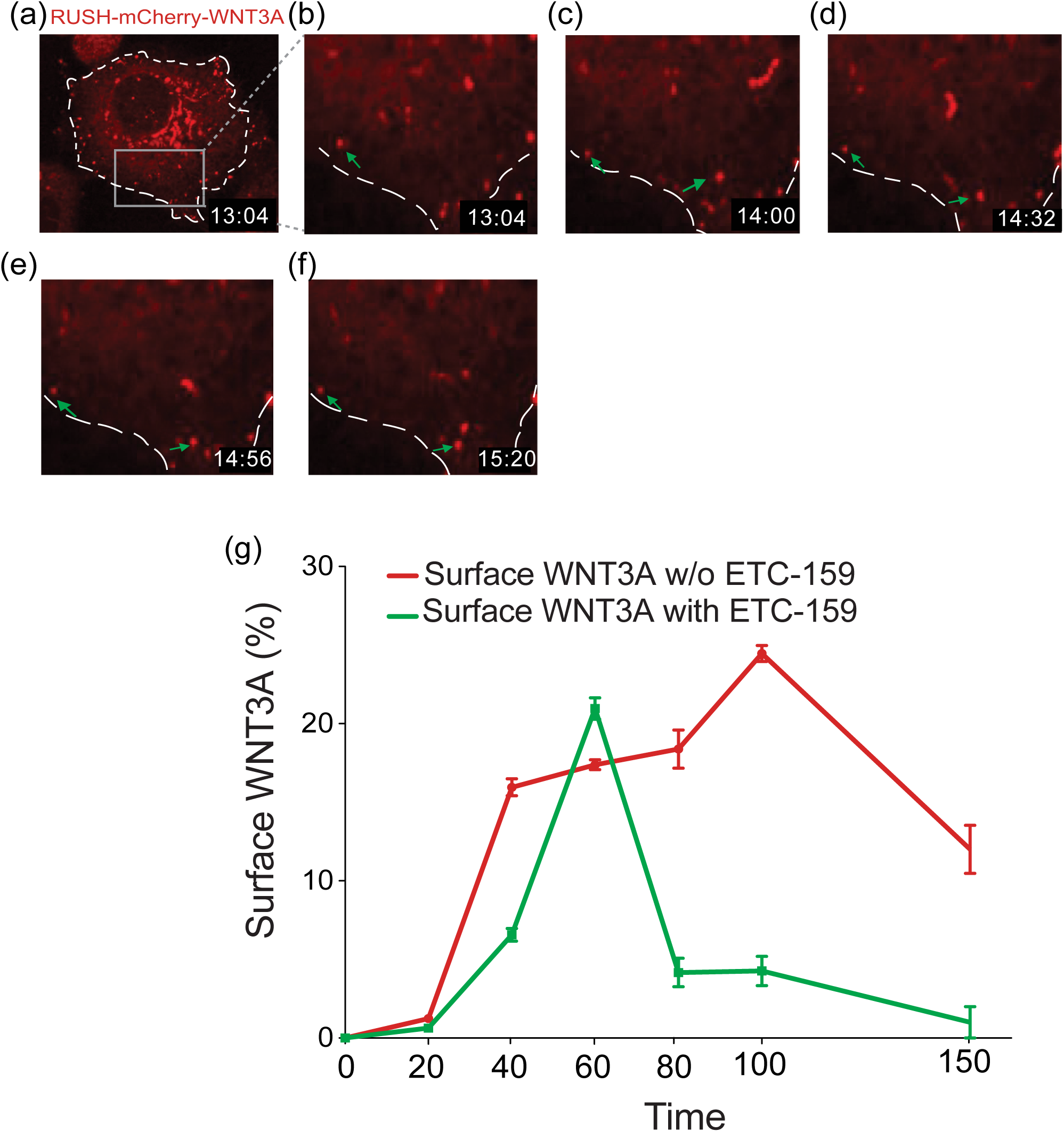
WNT3A time course to the membrane. Z-stack imaging reveals WNT3A vesicles lining up either inside or along the plasma membrane surface before moving out. Micrographs of z-stack images of HeLa cells expressing RUSH-WNT3A at the indicated time points after biotin addition. (a) Single cell. (b) magnification of indicated region. At 13:04, WNT3A started to move out of Golgi and aligning itself at sub membranous region. Inset shows WNT3A pausing at different regions (b-f). White arrow showing WNT3A moving to the plasma membrane whereas green showing WNT3A staying inside the cells. (g) Pulse-chase study of surface WNT3A on HeLa cells transfected with RUSH-WNT3A. Biotin and ETC-159 (100 nM) were added together where indicated. (average from n=3 independent FACS experiments).

How Wnts move from one cell to the next after secretion is an area of active study. We asked if Wnts remained bound to the cell surface of the producing cells or were rapidly secreted. Using antibodies to mCherry-WNT3A and flow cytometry in non-permeabilized cells, we studied the amount of time after biotin addition required for mCherry-WNT3A to reach the outside of the cell (Figure 4g). After washing and immunostaining, WNT3A was detectable at the cell surface 40 minutes after biotin addition and remained detectable there for the duration of the study period. However, the mCherry-WNT3A did not appear to accumulate over time, suggesting the Wnt was removed from the cell surface by release or re-endocytosis. To assess this, we performed a pulse-chase experiment. The PORCN inhibitor ETC-159 was added to the media at the same time as biotin, blocking the palmitoleation of newly synthesized WNT3A, so that only previously synthesized and palmitoleated WNT3A (the pulse) would be released and ‘chased’ from the ER. As the green line in Figure 4g illustrates, the pulse of newly released Wnt molecules are only transiently present on the cell surface.

### Wnts transfer from cell to cell via actin-based membrane extensions

Membrane extensions or protrusions, including filopodia and signaling filopodia (also called cytonemes), are important in cell migration, environmental sensing, and in generating mechanical forces. They also play a role in trafficking of membrane-bound molecules and signaling vesicles (Jacquemet et al. 2015; Leemhuis et al. 2010; Kornberg 2017). Limited studies on Wnt trafficking have shown that WNT8A can be transported on cytonemes from one cell to another (Stanganello et al. 2015). We used the RUSH-WNT3A model to further study Wnt trafficking via these membrane extensions. In a number of experiments, we were able to visualize RUSH-Wnt vesicles associated with long narrow cellular extensions. In Figure 5a, two adjacent HeLa cells expressing RUSH-Wnt3a passed Wnt from one cell to the other via a ∼1 um bridge, consistent with a filopodia (Supplementary Video S5A). These data further confirm Wnt trafficking along the filopodia extensions.

**Figure 5:**
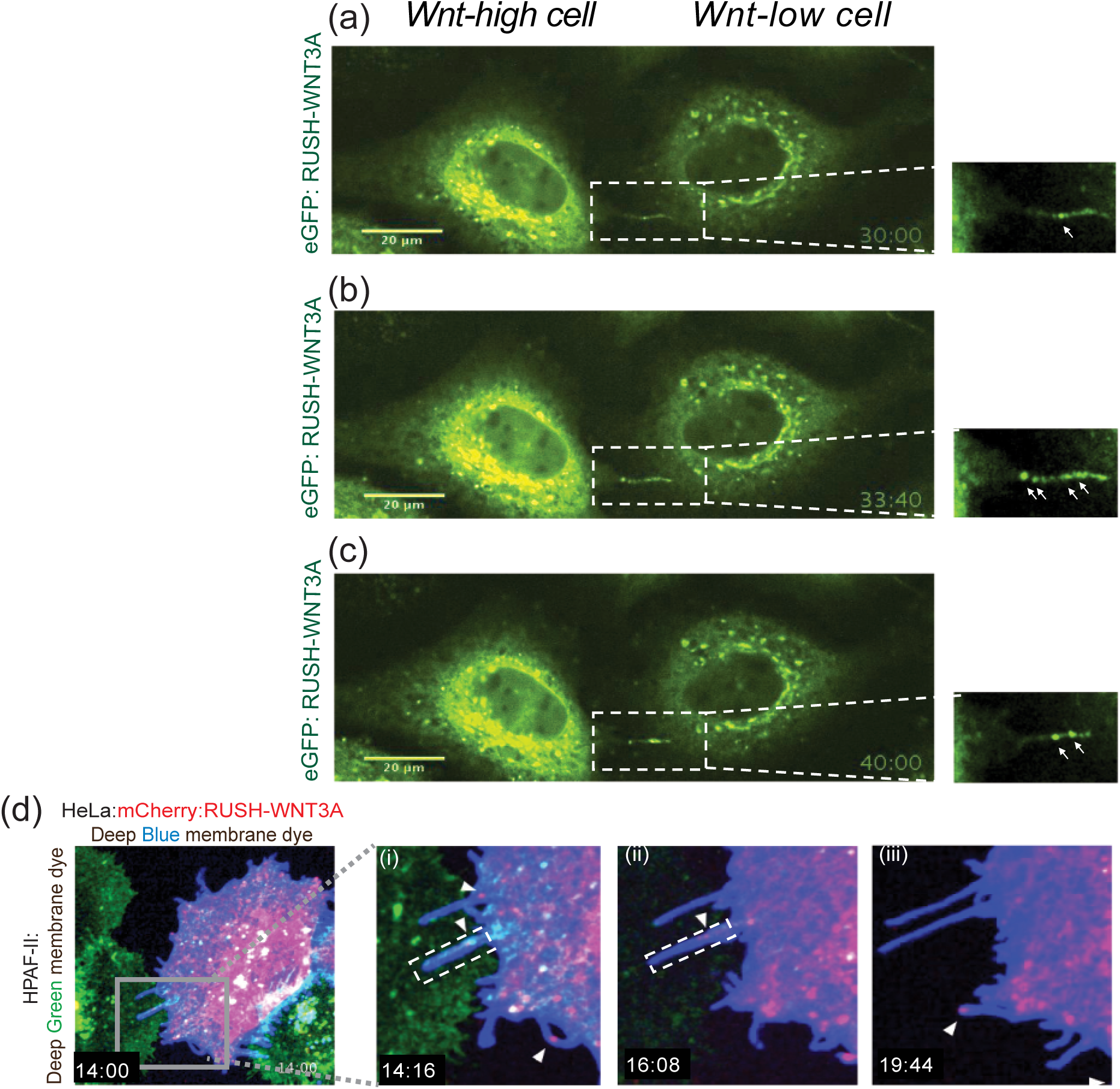
WNT3A transfer via filopodia. (a-c) Micrographs of adjacent HeLa cells expressing RUSH-WNT3A at indicated time points after biotin addition. From Supplemental video 5A. (a) WNT3A is in the Golgi after biotin treatment. Inset showing WNT3A movement along filopodia from one WNT3A-high cell to another WNT3A-low cell. Arrow shows filopodia extensions transporting WNT3A. (b) At t=33:40, multiple vesicles are seen with the filopodia. (c) At t=40:00. (d) Micrographs of HeLa cells (expressing RUSH-WNT3A(mCherry) and stained with CellMask™ Deep Blue membrane dye co-plated with HPAF-II cells stained with CellMask™ Deep Green membrane dye . From Supplemental Video 5B (i) Inset showing filopodia extending from HeLa cell to HPAF-II cell at 13:10. (ii-iii) Inset showing WNT3A (mCherry) on the filopodia from HeLa cells moving to HPAF-II cells at indicated time points.

Next, we co-plated RUSH-WNT3A expressing HeLa cells with HPAF-II cells to see if we could visualize Wnt transported from a Wnt-producing cell to a Wnt receiving cell. Upon biotin addition, RUSH-WNT3A from HeLa cells transported via apparent filopodia (Figure 5d, Supplementary Video S5B). The filopodia were typically between 1-6 μm in length although our system was not optimized to visualize longer extensions in multiple planes. Time-course z-stack confocal imaging revealed that RUSH-WNT3A (mCherry) can be seen transporting from HeLa cells to HPAF-II cells efficiently as compared to when HeLa cells plated alone (Supplementary Video S5B. These data suggest that WNT3A are transported from one cell to another via membrane protrusions resembling filopodia.

### LGR5-induced filopodia as Wnt signaling platforms

We wished to test if increasing the number of filopodia would increase the ability of a Wnt-producing cell to send a signaling to a neighboring Wnt-reporter cell. LGR5 is a Wnt target gene that can serve as an R-spondin (RSPO) co-receptor on Wnt receiving cells. However, more recent work has called this model into question (Lebensohn & Rohatgi 2018). Furthermore, LGR5 is constitutively internalized, a process that can be blocked by C-terminal tail mutations or by truncation at position 834 (834DEL). Forced retention of LGR5 at the cell surface induces a marked increase in the number of signaling filopodia on cells, (Carmon et al. 2012; Snyder et al. 2013; Snyder et al. 2015). We confirmed that co-expression of RUSH-mCherry-WNT3A and LGR5(834DEL)-eGFP led to a marked increase in the number of filopodia that retained trafficking of WNT3A vesicles (Figure 6a; Supplementary Video S6). STF assay further showed that LGR5-induced filopodia indeed increased the Wnt/ß-catenin signaling when cultured with HEK293 STF cells (Figure 6b). Consistent with the filopodia being actin-based, the number of filopodia and signaling were reduced when cells were cultured in the presence of very low dose latrunculin B (Figure 6b, c). This data is consistent with filopodia transmitting the Wnt signal from producing to receiving cell.

**Figure 6:**
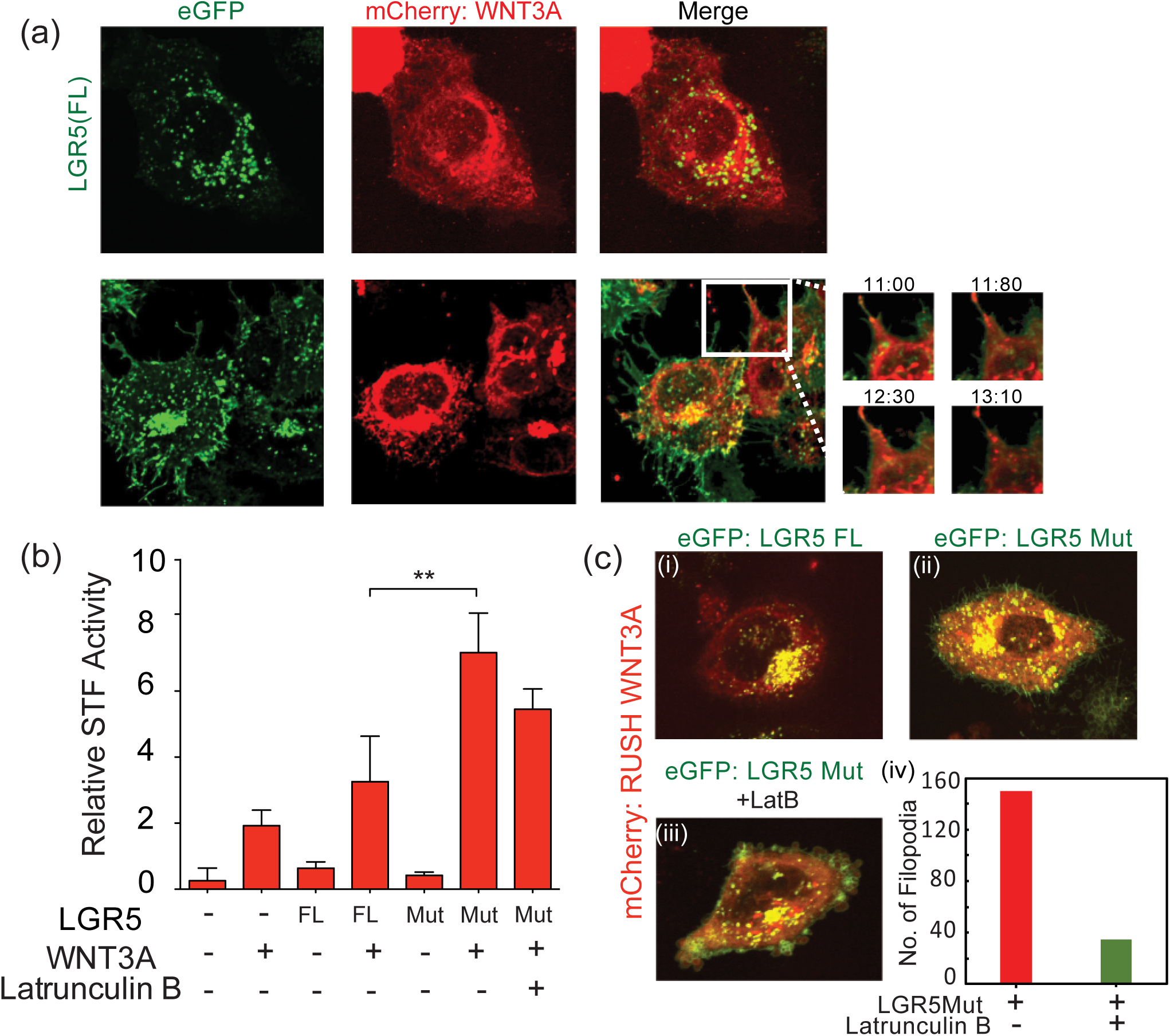
LGR5-induced filopodia as Wnt signalling conduits. (a) Z-stack micrographs of HeLa cells co-expressing RUSH-mCherry-WNT3A and eGFP-LGR5 full length (LGR_FL_) or LGR5_834DEL_, showing an increase in the number of filopodia in LGR5_834DEL_ mutant cells. Insets show movement of RUSH-WNT3A (mCherry) along the eGFP-LGR5-induced filopodia at different time points. (b) SuperTopFlash assay on HEK293 cells showing increased Wnt signaling in LGR5_834DEL_ mutant cells. Low dose latrunculin B was used to induce actin inhibition which shows slight decrease in Wnt signaling in LGR5_834DEL_ mutant cells. (\*\**p*<0.01) (c) Micrographs of HeLa cells transfected with LGR5 full length (LGR_FL_) (i) or LGR5_834DEL_ mutant cells (ii) showing increase in the number of filopodia that decreased upon treatment with latrunculin B (iv). (n=4).

## Discussion

Cell-to-cell communication is a vital step in the regulation of growth and development via secretory proteins. Understanding the mechanism underlying the transmission of signals through tissues is of profound interest and requires robust assays for their study. We show here that the quantitative and real-time RUSH system is well suited to study the production and movement of biologically active fluorescent-tagged Wnt proteins from the time of their initial synthesis to their movement to the cell surface (Augustin et al. 2012). The RUSH system also demonstrated Wnt-containing vesicles travelling via filopodia to neighboring cells, a phenomenon previously described in Drosophila and zebrafish but not to our knowledge in human cells. Supporting the physiological importance of Wnt filopodial transport, increasing the number of filopodia was able to increase Wnt/β-catenin signaling.

Initial studies on WLS showed it was a carrier for Wnts, but because those studies used a carboxyl-terminal epitope tag that masked the WLS ER localization signal, they mistakenly concluded that Wnt first binds to WLS in the Golgi. Subsequent work concluded that WLS binds Wnts in the ER, and here we confirm that in the absence of WLS, Wnt cannot exit the ER. Not surprisingly, in the presence of a PORCN inhibitor, Wnt that is not palmitoleated and hence cannot bind to WLS also cannot exit the ER. Thus, WLS appears to be the essential carrier of Wnts from the moment of its acylation to the plasma membrane, and likely, beyond (Gross et al. 2012; Korkut et al. 2009; Augustin et al. 2012).

One key finding here is the fate of Wnt-containing vesicles as they approach the plasma membrane. Live cell imaging shows these vesicles to pause near the plasma membrane prior to their disappearance. The data here support two fates for these vesicles – fusion with the membrane with release of Wnts to the outside of the cell, and more interestingly, passage via cellular extensions consistent with filopodia. These filopodia show active movement of Wnts, presumably in vesicles, towards neighboring cells. In other settings, these signaling filopodia are able to transmit signaling via signaling synapses. Further refinement of the RUSH system may facilitate the detection of Wnt signaling synapses in human cells and tissues. The concept of Wnt signaling synapses may be relevant in the mammalian intestine where Wnts are known to be transported across basement membranes, from the intestinal PDGFRα-expressing myofibroblasts to the stem cell compartment including the crypt base columnar (CBC) cells (Greicius et al. 2018) . The presence of extensive filopodial networks underlying the intestinal basement membrane has been noted by others (Cretoiu et al. 2017; Vannucchi et al. 2013).

LGR5 is a Wnt target gene expressed in the CBC cells and itself promotes Wnt/ß-catenin signaling (Barker et al. 2007; Chen et al. 2013; Sato et al. 2009). While LGR5 has been proposed as an R-Spondin receptor, Snyder et. al recently demonstrated, as we confirmed here, that an endocytosis-defective mutant of LGR5 stimulates the formation of filopodia and at the same time increased basal Wnt signaling. This is consistent with Wnt signaling utilizing filopodia even in monoculture. In addition, because LGR5 is a Wnt target gene, it suggests the possibility of a positive Wnt feedback loop, where Wnt signaling from the stroma to the CBC cells stimulates LGR5-driven production of receiving filopodia. Since not all Wnt-responsive cells in the intestinal crypt produce LGR5, this could be a mechanism to lock stem cells into a circuit with the Wnt producing cells while providing more graded amount of Wnt signaling to LGR5(−) cells.

Previous studies show that the active Wnt signaling occurs at plasma membrane (PM). It is of interest to examine both the Wnt-producing and Wnt-receiving cells. Our data highlights an important demarcation between the two cell types. Our data suggest Wnts do not reside for long on the PM of Wnt-producing cell. The newly produced Wnt may be re-endocytosed (Pfeiffer et al., 2002) or passed to the Wnt-receiving cell where it forms a signalosome (Bilic et al., 2007) and initiates a Wnt/ß-catenin dependent signaling event (Li et al. 2012; Posokhova et al. 2014)(Li et al., 2012; Posokhova et al., 2015). Our data suggests a method of WNT3A transport from Wnt-producing cells, where Wnt vesicles pause near subcellular junctions of PM. A subset of these vesicles either fuse with the PM, or are passed to Wnt-receiving cells via cellular extensions resembling actin-based filopodia. Our images suggest that these filopodia vary in length. HPAF-II cells that were close to Wnt-producing HeLa cells showed the longest nanotubes with WNT3A on it (Figure 5d). There are instances where WNT3A-containing nanotubes retracts back and we assume this is due to absence of a nearby target cell. These data are consistent with filopodia as a transport mechanism for WNT3A.

The RUSH-Wnt system is a useful model system to study Wnt transport. It is relatively straightforward to utilize in cultured cells. The RUSH-Wnt system may also be useful in high-content screening applications. Further refinement may allow the study of Wnt transport in more complex systems such as organoid cultures to provide insights into how Wnts move from stroma Wnt producing cells to tissue stem cells.

## Acknowledgments

We acknowledge the assistance of members of the Virshup lab, and Frederic Bard for fruitful discussion, and Babita Madan for advice and critical reading of the manuscript. We also acknowledge the assistance of SingHealth Advanced Bioimaging core, specially Wong Wei Juan and Keshmarathy Sacadevan for their assistance with microscope and image analysis software. The research is supported by Singapore Ministry of Health’s National Medical Research Council under the STAR Award Program to DMV.

## Supplemental Figures Legends

**Supplemental Figure S1: Synchronization of WNT8A protein using RUSH system.**

(a) Indirect immunofluorescence photomicrographs of HeLa cells expressing RUSH-eGFP-WNT8A at various time points after biotin addition, from Supplemental Video S1B. At time 00:00 (minutes:seconds) of biotin addition, WNT8A remained in the ER. At time 10:00, WNT8A can be seen in the Golgi. At time 20:00, WNT8A is seen entirely in Golgi. Starting around 45:00, WNT8A can be seen in vesicle exiting the Golgi and moving towards PM. By time 54:00, the Golgi begins to be depleted of WNT8A.

(b) Time-dependent analysis of ER-Golgi localization of RUSH-WNT8A. The plot shows fluorescence intensity in the Golgi region (white box in Figure (a)) at different time point after biotin addition. Intensities were normalized to maximum Golgi intensity. (n=5 cells)

## Supplemental Videos Legends

**Supplemental Video S1A:** Real-time imaging of the synchronized trafficking of RUSH-eGFP-WNT3A (corresponds to Fig. 1). HeLa cells were transfected to express KDEL-Streptavidin as a hook and SBP-eGFP-WNT3A as a reporter. After 18 h of expression, at time 00:00, 100 μM biotin was added to induce the release and monitored using Nikon’s spinning disk confocal microscope.

**Supplemental Video S1B:** Real-time imaging of the synchronized trafficking of RUSH-WNT8A (corresponds to Supp Fig. S1). HeLa cells were transfected to express KDEL-Streptavidin as a hook and SBP-eGFP-WNT8A as a reporter. After 18 h of expression, at time 00:00, 100 μM biotin was added to induce the release and monitored using Nikon’s spinning disk confocal microscope.

**Supplemental Video S2:** Real-time imaging of the synchronized trafficking of RUSH-WNT3A in the presence and absence of known PORCN inhibitor, ETC-159 (corresponds to Fig. 2). HeLa cells were transfected with RUSH-WNT3A and after 6-7 h of transfection, treated with ETC-159. 100 μM biotin was added ∼12 h later.

**Supplemental Video S3A:** Real-time imaging of the synchronized trafficking of RUSH-WNT3A in RKO WT and RKO WLS KO cells (corresponds to Fig. 3). Cells were transfected with RUSH-eGFP-WNT3A plasmid and 100 μM biotin was added 18 h later.

**Supplemental Video S3B:** Real-time imaging of the synchronized trafficking of RUSH-WNT3A with and without exogenous WLS. RKO WLS KO cells were transfected with RUSH-mCherry-WNT3A plasmid and 100 μM biotin was added 18 h later.

**Supplemental Video S4:** Real-time z-stack imaging of the synchronized trafficking of RUSH-WNT3A (corresponds to Fig. 4). HeLa cells were transfected with RUSH-mCherry-WNT3A plasmid and after 18 h of expression, 100 μM biotin was added and monitored using Nikon’s spinning disk confocal microscope. Z-stacks were analysed and merged on Fiji 2.0. Image acquisition was started ∼12 min after biotin addition to minimize photo bleaching.

**Supplemental Video S5A:** WNT3A transfer via filopodia. Real-time imaging of the synchronized trafficking of RUSH-WNT3A (corresponds to Fig. 5a). HeLa cells were transfected with RUSH-mCherry-WNT3A plasmid and after 18 h of expression, 100 μM biotin was added and monitored using Nikon’s spinning disk confocal microscope.

**Supplemental Video S5B:** Co-culture of Wnt producing and Wnt receiving cells. Real-time imaging of the synchronized trafficking of RUSH-WNT3A (corresponds to Fig. 5b). HeLa cells transfected with RUSH-WNT3A and stained with CellMask™ Deep Blue membrane dye were co-plated with HPAF-II cells stained with CellMask™ Deep Green membrane dye. After 18 h of expression, 100 μM biotin was added and monitored using Nikon’s spinning disk confocal microscope. Images were acquired ∼12 minutes after biotin addition to minimize photobleaching.

**Supplemental Video S6:** Mutant LGR5-induced filopodia transmit Wnt vesicles. Real-time imaging of the synchronized trafficking of RUSH-WNT3A (corresponds to Fig. 6a). HeLa cells co-transfected with RUSH-mCherry-WNT3A and eGFP-LGR5 834DEL mutant plasmids. After 18 h of expression, 100 μM biotin was added and monitored using Nikon’s spinning disk confocal microscope. Images were acquired ∼12 minutes after biotin addition to minimize photobleaching.

